# Deep structural brain lesions associated with consciousness impairment early after haemorrhagic stroke

**DOI:** 10.1101/441709

**Authors:** Benjamin Rohaut, Kevin W. Doyle, Alexandra S. Reynolds, Kay Igwe, Caroline Couch, Adu Matory, Batool Rizvi, David Roh, Angela Velasquez, Murad Megjhani, Soojin Park, Sachin Agarwal, Christine M. Mauro, Gen Li, Andrey Eliseyev, Vincent Perlbarg, E. Sander Connolly, Adam M. Brickman, Jan Claassen

**Author notes:** **Address for Correspondence and Reprints**, Jan Claassen, MD, PhD, Neurological Institute, Columbia University, 177 Fort Washington Avenue, MHB 8 Center, Room 300 New York, NY 10032. **Financial Disclosure Statement:** All authors declare no financial relationships with any organisations that might have an interest in the submitted work and, no other relationships or activities that could appear to have influenced the submitted work. The Corresponding Author affirms that the manuscript is an honest, accurate, and transparent account of the study being reported; that no important aspects of the study have been omitted; and that any discrepancies from the study as planned have been explained.

## Abstract

**Background:** The significance of deep structural lesions on level of consciousness early after intracerebral haemorrhage (ICH) is largely unknown.

**Methods:** We studied a consecutive series of patients with spontaneous ICH that underwent MRI within 7 days of the bleed. We assessed consciousness by testing for command following from time of MRI to hospital discharge, and determined 3-months functional outcomes using the Glasgow Outcome Scale-Extended (GOS-E). ICH and oedema volumes, intraventricular haemorrhage (IVH), and midline shift (MLS) were quantified. Presence of blood and oedema in deep brain regions previously implicated in consciousness were assessed. A machine learning approach using logistic regression with elastic net regularization was applied to identify parameters that best predicted consciousness at discharge controlling for confounders.

**Results:** From 158 ICH patients that underwent MRI, 66% (N=105) were conscious and 34% (N=53) unconscious at the time of MRI. Almost half of unconscious patients (49%, N= 26) recovered consciousness by ICU discharge. Focal lesions within subcortical structures predicted persistent impairment of consciousness at discharge together with MLS, IVH, and ICH and oedema volumes (AUC 0.74; 95%-CI 0.73-0.75). Caudate nucleus, midbrain peduncle, and pontine tegmentum were implicated as critical structures. Unconscious patients predicted to recover consciousness had better 3-month functional outcomes than those predicted to remain unconscious (35% vs 0% GOS-E ≥4; p-value=0.02).

**Conclusion:** MRI lesions within key subcortical structures together with measures reflecting the mass effect of the haemorrhage (lesion volumes, IVH, MLS) obtained within one week of ICH can help predict early recovery of consciousness and 3-month functional outcome.

## Introduction

Insights into mechanisms underlying early and delayed recovery of consciousness following brain injury are limited. Investigators have implicated several subcortical structures to be crucial for maintenance of arousal such as pontine tegmentum, midbrain, basal forebrain, hypothalamus and central thalamus [1–3]. Structures important for conscious processing or awareness include sub-cortical regions (i.e., thalamus, putamen, caudate and pallidum) as well as associative cortical regions (i.e., prefrontal, temporal and parietal cortices) and their connecting neuronal pathways [4–9]. Based on neuropathological, imaging and electrophysiological studies, circuit models have been developed that help conceptualize impairment and recovery of consciousness (e.g., modern concepts of the ascending reticular activating pathway [ARAS],[3,10,11] mesocircuit model,[6,12] and global neuronal workspace[5]). Structural damage to regions within this circuitry (e.g., large intracerebral haemorrhage [ICH] in the thalamus) as well as lesions that functionally affect circuit connectivity (e.g., stretching thalamo-cortical projections from mass effect), especially when they are bilateral, may result in clinically indistinguishable unconscious patients. However, depending on the anatomical location and the pathophysiological mechanism of lesions, prognosis for recovery of consciousness can dramatically differ [13].

Here we studied patients with ICH, a condition that may cause both focal injury to specific subcortical brain regions that are integral parts of the ARAS and/or the mesocircuit model as well as more diffuse injury that can impair network connectivity (e.g., from midline shift and/or oedema) [14]. Specifically, we explored how the level of impairment and recovery of consciousness relate to the locations and the extent of subcortical injury quantified by early MRI. We tested the hypothesis that focal lesions within subcortical regions included in the previously mentioned models of consciousness, in addition to established characteristics of the haemorrhage (i.e., volume of ICH and oedema, and midline shift [MLS]), contribute to consciousness level during the acute phase of ICH.

## Methods

### Subjects

We studied a consecutive series of patients with ICH that underwent MRI including fluid attenuated inversion recovery (FLAIR) and diffusion weighted imaging (DWI) within one week of the ICH between March 2009 and November 2015. Inclusion criteria were: (1) spontaneous ICH, and (2) MRI obtained within 7 days of the haemorrhage. Exclusion criteria were: (1) age <18 years, (2) pregnancy, (3) ICH due to tumour, trauma, or haemorrhagic conversion of an ischemic stroke, and (4) patients or families who declined to participate in the study. Patient management was in accordance with current guidelines (Supplementary Material). Data were collected as part of a prospective observational cohort study approved by the local institutional review board. Written informed consent was obtained from patients and/or legal surrogates.

### Clinical variables

We collected baseline demographic and medical history (e.g., age, gender, race), and admission characteristics of the ICH (e.g., ICH volume and location, presumed aetiology, intraventricular haemorrhage, primary ICH-score)^8^. We calculated the admission Functional Outcome in Patients With Primary Intracerebral Hemorrhage (FUNC) score by quantifying ICH volume and location, age, Glasgow Coma Scale, and pre-ICH cognitive impairment.[16] Daily assessments included documentation of seizures (as per hospital protocol all unconscious patients undergo continuous EEG monitoring for at least 24 hours), metabolic abnormalities (e.g., renal function and liver failure), and fever. Doses of all sedatives and laboratory values were recorded at the time of all behavioural assessments.

### Behavioural assessment

We assessed level of consciousness daily from ICU admission to 30 days post-injury or hospital discharge, whichever was sooner. As described previously [17], behavioural assessments of consciousness were performed during morning rounds. These consisted of protocolized, hierarchical assessments categorizing consciousness into three levels of behavioural states: (1) “comatose” (no response to stimulation), (2) “arousable” (opening eyes and/or attending to stimulation), or (3) “conscious” (following simple commands; e.g., “show me two fingers”). To overcome language impairment or aphasia while testing for consciousness, we used in addition to verbal commands, non-verbal cues to induce mimicking (e.g., holding up two fingers and then gesturing to subject’s supported hand). For the classification approach described below, we dichotomized patients into “conscious” (category 3, following verbal and/or non-verbal commands) and “unconscious” (categories 1 and 2; see details in supplementary Material). According to our ICU protocol daily assessments were performed during interruption of sedation.

### MR acquisition

As part of our clinical protocol we acquired MR images within 7 days of haemorrhage whenever deemed safe by the attending neurointensivist using a 3T scanner (GE Signa HDx MRI scanner; HD23 software). Total acquisition time did not exceed 45 minutes. We obtained FLAIR, T1-weighted, and DWI sequences (for details please refer to the supplementary material section).

### Categorization of lesions

Anatomical regions of interest (ROIs) were predefined based on established neuroanatomical atlases [18] with a focus on subcortical brain regions (henceforth referred to as “subcortical ROIs”) previously implicated in consciousness [1–7, 12]. A board-certified neurologist (AR) categorized the presence of blood and perihematomal FLAIR hyperintensity (henceforth referred to as “oedema”) for each ROI based on a 3D visualization of FLAIR, T1 and DWI sequences. The following ROIs were included in the models: pontine tegmentum, midbrain (central and peduncles), hypothalamus, basal forebrain, thalamus, pallidum, putamen, and caudate nuclei (see Figure 1, 1^st^ row in pink colour). For purposes of analysis, lesion laterality was reclassified from right/left into ipsi/contralateral using the following approach. The side of the brain with the larger amount of blood was labelled as ipsilateral. The side with the smaller amount was labelled as contralateral. Intraventricular haemorrhage (IVH) was assessed in the 3^rd^, 4^th^, and each lateral ventricle and classified as present or absent. Any challenging cases with bilateral haemorrhage were classified by consensus between three board certified neurologists (AR, JC, BR). A board-certified neurologist (DR) coded the same imaging parameters on a random 20% sample of MRIs blinded to the first coder’s results. Interrater agreement was assessed using kappa statistics.

**Figure 1.**
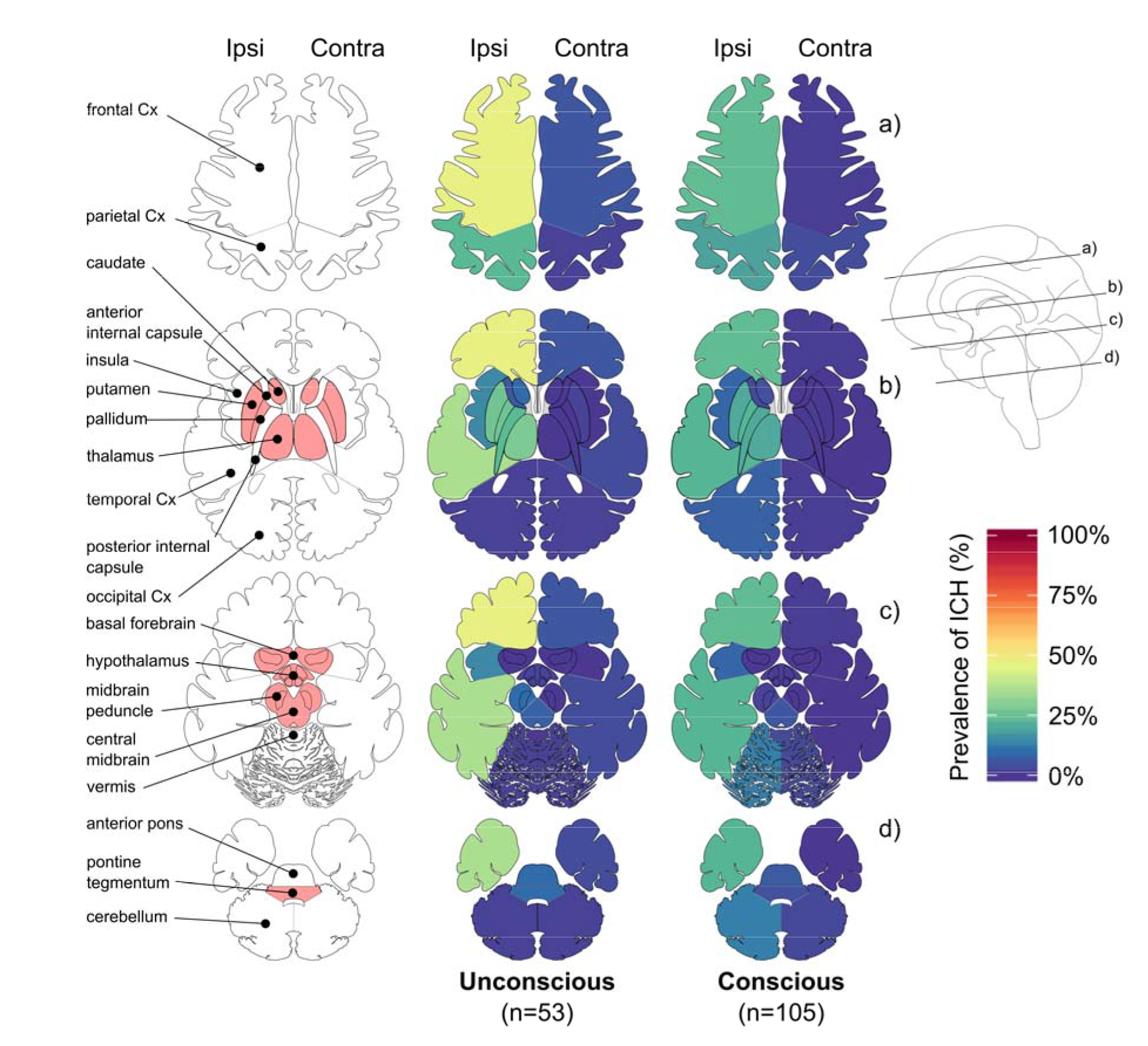
Anatomical segmentation, subcortical ROIs and location of haemorrhages according to consciousness level. Anatomical segmentation is represented on the left pane with the explored subcortical regions of interest (ROIs) in pink. Prevalence of ICH observed on MRI are showed by level of consciousness at the time of the MRI (“unconscious”: patients did not follow or mimic even simple commands; “conscious”: patients followed or mimicked simple commands. “ipsi” and “contra” stand for ipsilateral and contralateral with respect to the primary side of the haemorrhage; Cx: cortex.

### Volumetric measurements and midline shift

Haemorrhage, perilesional oedema, and brain volumes were quantified based on FLAIR sequences using a semi-automatized method. Briefly, a gross region-of-interest was identified that encapsulated the affected region (ICH or oedema) to automatically compute a 3D image that were visually inspected and manually corrected if necessary (KI, see Supplementary Material and Figure 2 panel A). Midline shift (MLS) was measured both at the level of the septum pellucidum as well as at the pineal gland, and the larger number was recorded [19].

### Main outcome

Main outcomes were the level of consciousness observed at time of ICU discharge and the Glasgow Outcome Scale-Extended (GOS-E) obtained 3 months following the haemorrhage via phone interviews [20]. As an additional outcome measure, we recorded the best level of consciousness observed at any time during hospitalization following MRI acquisition.

### Confounders

All patients were clinically evaluated for the presence of seizures, hypo-or hyperglycaemia (70 and 200 mg/dL, respectively), hypo-and hypernatremia (133 and 150 mmol/L, respectively), and renal and fulminant liver failure at the time of behavioural assessments. All analyses were directly controlled for potential metabolic confounders (including blood urea nitrogen, creatinine, serum glucose level). In addition to the above outlined protocol of stopping sedation for all behavioural assessments we collected the cumulative doses of all sedative medications administered within the two elimination half-lives preceding clinical assessments.[17].

**Figure 2.**
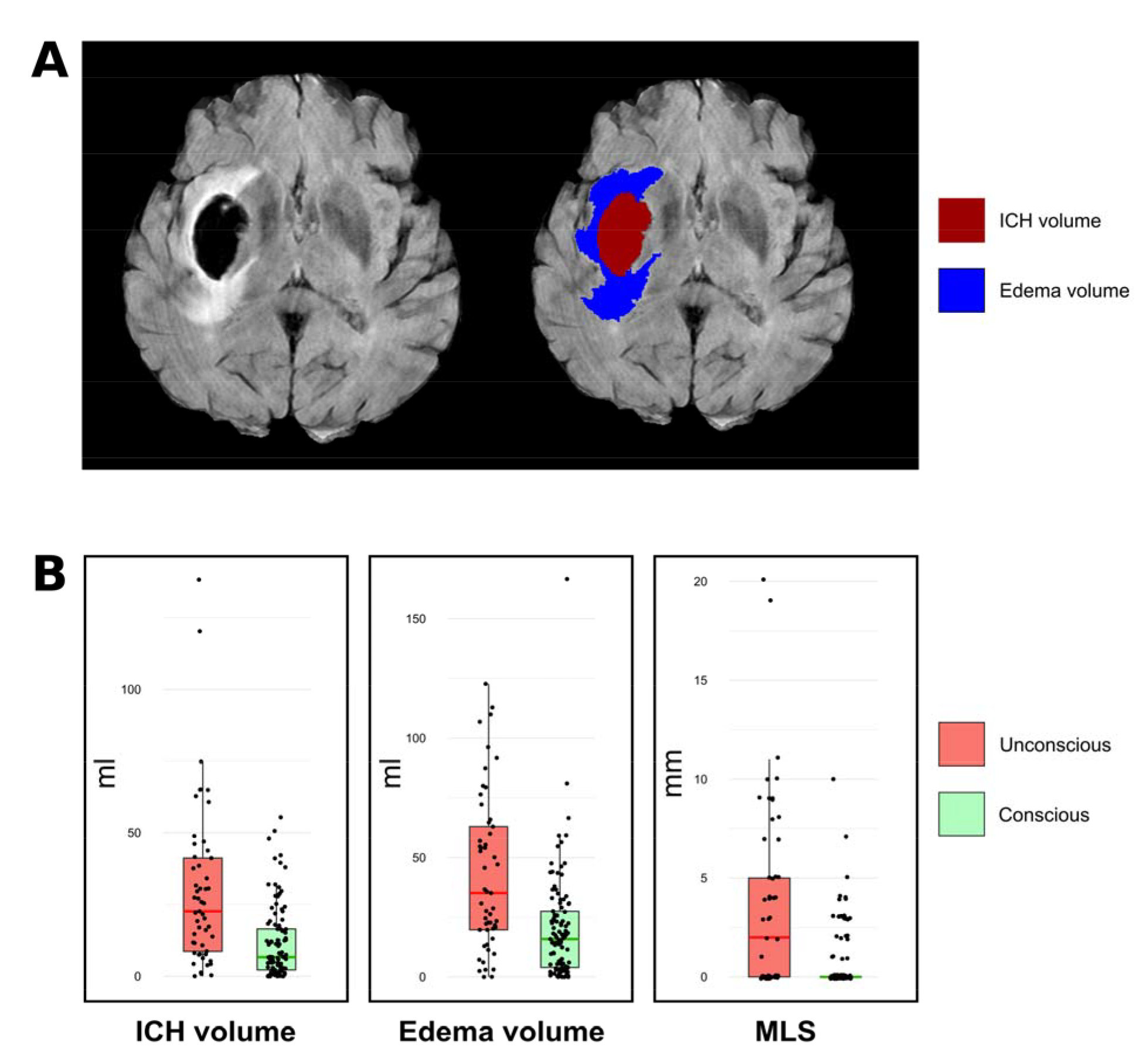
Haemorrhage and oedema volumes and midline shift. Panel A. Illustrates the volume on MRIs of one exemplary case. **Panel B.** Measurements according to consciousness level at time of MRI (normalized values; for details please refer to methods). ICH: Intracerebral Hemorrhage; MLS: midline shift.

### Statistical analysis

A machine learning approach using logistic regressions with elastic net regularization was applied to identify the parameters that best predicted consciousness at time of MRI and at time of ICU discharge [21]. This method allows a robust data-driven analysis when there are a large number of features compared to the number of observed events and/or when features are highly correlated. Models were trained on the clinical labels (conscious vs unconscious) obtained either at the time of MRI or at the time of ICU discharge. In order asses robustness of the model, we performed 5-fold cross validations repeated 500 times [22]. Model performance was evaluated using the area under the receiver operating characteristic curve (AUC) with 95% confidence intervals (95% CI). Logistic regression using elastic net regularization were computed with the Glmnet R package (for details please refer to the Supplementary Material).

Differences in baseline features between the patients that fulfilled inclusion criteria and those that did not were explored using Fisher’s exact test for categorical and Wilcoxon–Mann–Whitney test for quantitative variables as appropriate. All statistical tests were two-sided. Categorical variables are reported as percentage (number) and quantitative variables as median (interquartile range). Significance was set at P<0.05. All analyses were performed using the R statistical software version 3.4.1 [23].

## Results

### Enrolment bias analysis

From a total of 690 patients admitted during the study period, 23% (N=158) had an MRI within the 7 days of the haemorrhage and fulfilled the inclusion criteria. Patients included in the study more frequently had presumed amyloid as the underlying aetiology, lobar location, better admission GCS, primary ICH and FUNC scores, smaller ICH volumes on the admission CT scan, and better outcomes as reflected in the 3-months GOS-E, when compared to patients that were not included (Table 1). MRI scans were obtained within a median of 2 (IQR 1, 3) days from ICH. Main ICH aetiologies were hypertension (49%; N=78) and amyloid (37%; N=59).

**Table 1.** Baseline characteristics of the reported cohort and enrolment bias.

|  | **MRI**<br>**(N=158)** | **No MRI**<br>**(N=532)** | **p-value** |
| --- | --- | --- | --- |
| Age, years | 68 [54, 77] | 63 [50, 76] | 0.05 |
| Female | 71 (45) | 244 (47) | 0.6 |
| White | 51 (32) | 148 (29) | 0.4 |
| Presumed aetiology |  |  | < 0.01 |
| Hypertensive | 78 (49) | 241 (48) |  |
| Amyloid | 59 (37) | 62 (12) |  |
| Coagulopathy* | 16 (10) | 86 (17) |  |
| Other | 5 (3) | 121 (24) |  |
| GCS at admission | 14 [9,15] | 10 [5, 15] | < 0.01 |
| **ICH characteristics on admission CT** |  |  |  |
| Lobar | 56 (35) | 110 (21) | < 0.01 |
| Deep | 62 (39) | 240 (47) | 0.1 |
| Infratentorial | 28 (18) | - | - |
| ICH vol |  |  | < 0.01 |
| <30 | 121 (78) | 306 (66) |  |
| 30-60 | 27 (17) | 89 (19) |  |
| >60 | 7 (5) | 69 (15) |  |
| IVH | 72 (47) | 281 (59)** | < 0.01 |
| **ICH prognostic scores** |  |  |  |
| ICH score | 1[1, 2] | 2 [1, 3] | < 0.01 |
| FUNC score | 9[7, 10] | 8 [6, 9] | < 0.01 |
| **Hospital course** |  |  |  |
| EVD | 31 (20) | 142 (27) | 0.05 |
| Clot evacuation | 16 (10) | 69 (13) | 0.3 |
| ICU stay, days | 4 [2, 8] | - |  |
| Hospital stay, days | 10 [6, 20] | - |  |
| **Outcome 3 months** |  |  |  |
| GOS | 3 [2, 4] | 3 [2, 5] | < 0.01 |
| GOS-E | 4 [1, 6] | - |  |
| Dead | 24 (22) | 158 (42)** | < 0.01 |

Data reported as n (%) or median [25%-IQR, 75%-IQR] as appropriate.
MRI: Magnetic Resonance Imaging; GCS: Glasgow Coma Scale; ICH: Intracerebral Haemorrhage; CT: Computed Tomography; IVH: Intraventricular Haemorrhage; EVD: External Ventricular Drain; ICU: Intensive Care Unit; GOS: Glasgow Outcome Scale; GOS-E: Glasgow Outcome Scale – Extended. \* Coagulopathy, primary haematological disorder and medication induced combined; \*\* more than 5% missing data.

### Patient cohort

From a total of 158 ICH patients, 66% (N=105) were conscious, and 34% (N=53) were unconscious at the time of MRI (Figure 3). At ICU discharge (occurring on median day 4 [2, 8]), 79% (N=125) were conscious, 18% (N=28) remained unconscious, and 3% were dead (N=5). 49% (N=26) of initially unconscious patients recovered consciousness at ICU discharge. 6% (N=6) of the initially conscious patients became unconscious during the ICU stay. Reasons for secondary unconsciousness included worsening oedema (N=3), hydrocephalus (N=1), ventriculitis (N=1), and seizures (N=1). At the time of MRI acquisition and ICU discharge, hypo-or hyperglycaemia, hypo-or hypernatremia, renal or fulminant liver failure were not present to explain unconsciousness. One patient with secondary loss of consciousness was seizing prior to death (for the purposes of the study this was considered the ICU discharge time).

**Figure 3.**
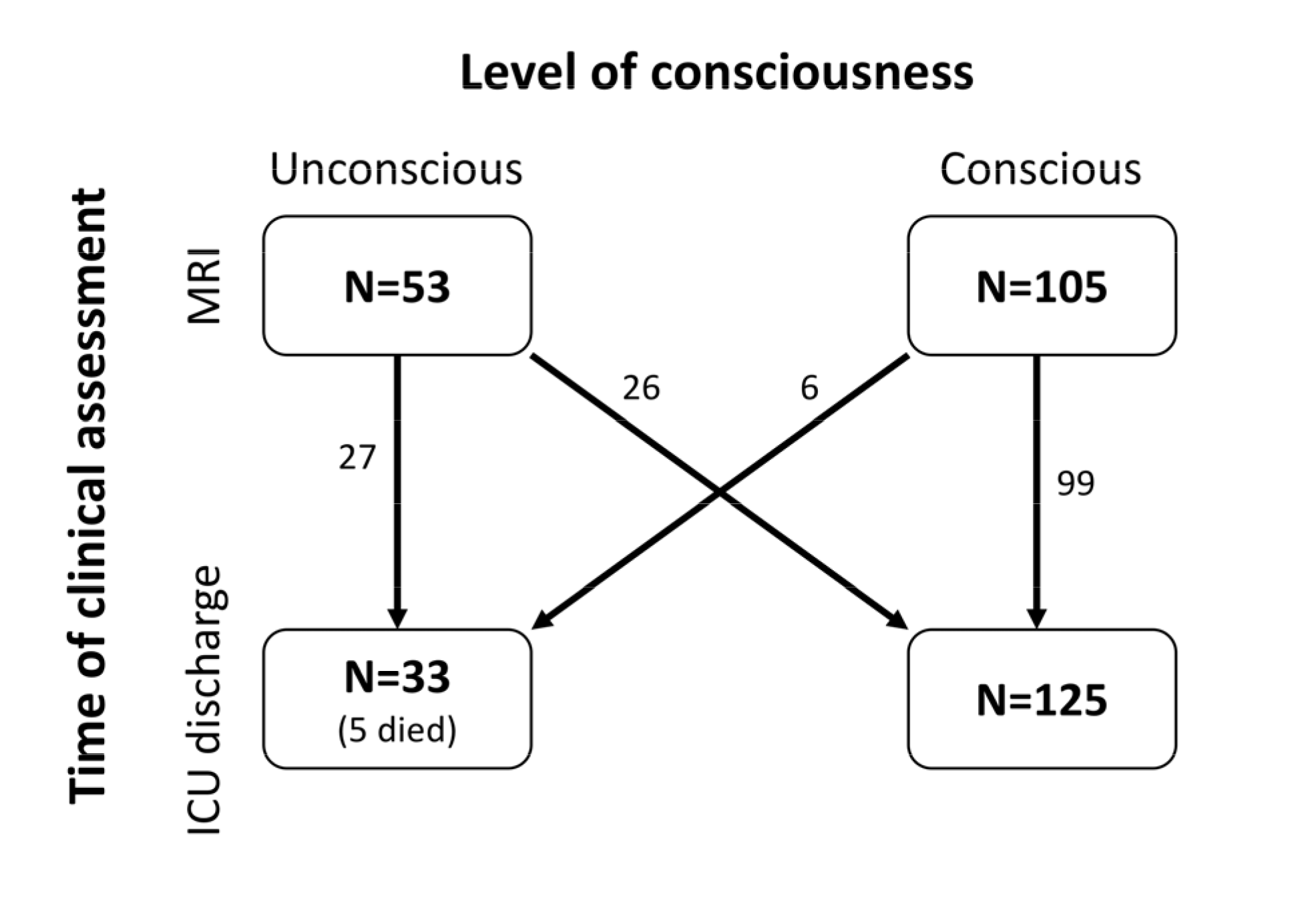
Flow chart. Level of consciousness assessed at MRI and ICU discharge. Note that for the 5 patients who died in the ICU, we considered the last neurological exam as the assessment at ICU discharge (of those that died, 3 patients were unconscious and 2 conscious at time of MRI, all of them were unconscious prior to death).

At time of MRI scan only 15% (N=24) of patients received any sedative medication (see table S1) and none of the patients received sedatives at ICU discharge. In patients that were unconscious at the time of MRI, propofol had been administered more frequently (19% vs 6%) but at lower cumulative doses (131+/-69 vs 361+/-202 mg over the 2 preceding half-lives) than those that were conscious.

### MRI findings

73% (N=116) of patients had an isolated unilateral supratentorial ICH whereas 11% (N=17) had an isolated infratentorial ICH and 7% (N=11) had both (supratentorial and infratentorial ICH). ICH was most frequently observed in frontal (37%) and temporal cortices (27%), globus pallidus (23%), thalamus (23%), putamen (22%), posterior limb of internal capsule (21%), and the parietal cortex (22%, Figure 4). Interrater agreements for both ICH and oedema measures in all ROIs reached a median kappa of 0.82 (0.66, 0.88).

Patients that were unconscious at the time of MRI more frequently had ICH affecting frontal and temporal lobes as well as the brainstem. Amongst unconscious patients, ICH was least frequently located in the cerebellum and occipital cortex. Brain volumes ranged from 1101 to 1887ml (group mean 1451ml). After normalization, ICH volumes were 11 ml (4, 26), oedema volumes 21 ml (8, 39), and MLS was 0 (0, 3) mm. ICH volume, oedema volume, and MLS were greater for patients that were unconscious at time of MRI and at time of ICU discharge when compared to those that were conscious (Figure 2; Table S2). IVH was more common amongst unconscious patients at ICU discharge (Table S2).

### Prediction of consciousness at time of MRI and ICU discharge

The algorithms trained on subcortical ROIs together with ICH and oedema volumes, MLS and IVH accurately predicted level of consciousness both at the time of MRI and at ICU discharge (AUC = 0.74 [95%CI: 0.72, 0.75] and 0.74 [95%CI: 0.73, 0.75], respectively). ICH volume and MLS were the most important predictors (Table 2). Lesions in pontine tegmentum (oedema), in the ipsilateral caudate nuclei (ICH and oedema), as well as in the contralateral midbrain peduncle, putamen and pallidum (oedema), were identified as predictors of being unconscious at time of MRI (Table 2; Figure S1). Lesions in pontine tegmentum (oedema), ipsilateral midbrain peduncle (ICH) and ipsilateral caudate nuclei (ICH and oedema; please refer to the Supplementary Material for details on these patients) were identified as predictors of being unconscious at ICU discharge (Table 2; Figure S2). In addition, over both models, a few ROIs were found to be associated with being conscious (e.g., pallidum, central midbrain, basal forebrain; see Table 2, Figure S1 and S2 and discussion).

**Figure 4.**
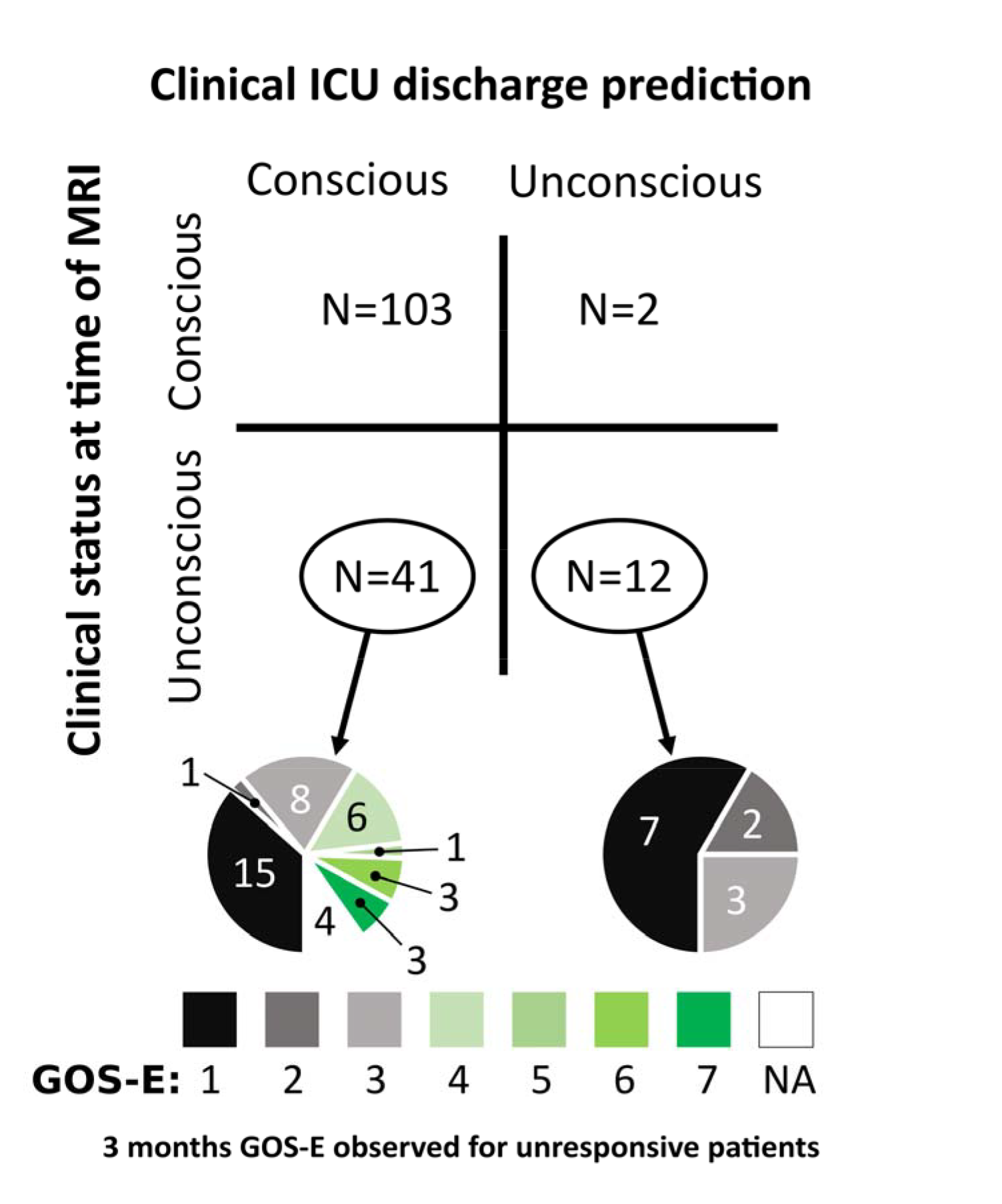
Predicted level of consciousness at ICU discharge based on imaging measures. Unconscious patients at time of MRI that were predicted to be conscious at ICU discharge (n=41) based on imaging findings were more likely to be conscious at ICU discharge and had a greater chance to reach a GOS-E ≥ 4 at 3 months (illustrated in shades of green; p-value = 0.02). GOS-E: Glasgow Outcome Scale – revised; NA: not available (lost follow-up).

**Table 2.** Weights of models predicting consciousness at time of MRI and at time of ICU discharge.

| Subcortical ROIs | Conscious at time of MRI |  | Conscious at time of ICU discharge |  |
| --- | --- | --- | --- | --- |
|  | ICH | Oedema | ICH | Oedema |
| Caudate ipsi | **-0.12[-0.7, 0]** | **-0.32[-0.65, 0]** | **-0.78[-1.14, -0.25]** | **-0.72[-0.99, -0.48]** |
| Caudate contra | 0[0, 0] | 0[0, 0] | 0[0, 0] | 0[0, 0] |
| Putamen ipsi | 0[0, 0] | 0[0, 0] | 0[0, 0] | 0[0, 0] |
| Putamen contra | **0[-0.03, 0]** | **0[-0.32, 0]** | 0[0, 0] | 0[0, 0] |
| Pallidum ipsi | 0[0, 0] | **0.31[0, 0.72]** | 0[0, 0] | 0[0, 0] |
| Pallidum contra | **0[0, 0.67]** | **0[-0.82, 0]** | 0[0, 0] | 0[0, 0] |
| Thalamus ipsi | **0[-0.06, 0]** | 0[0, 0] | 0[0, 0] | 0[0, 0] |
| Thalamus contra | 0[0, 0] | 0[0, 0] | 0[0, 0] | 0[0, 0] |
| Basal forebrain | 0[0, 0] | 0[0, 0] | 0[0, 0] | **0[0, 0.42]** |
| Hypothalamus | 0[0, 0] | **0[-0.06, 0]** | 0[0, 0] | 0[0, 0] |
| Midbrain peduncle ipsi | 0[0, 0] | 0[0, 0] | **0[-0.12, 0]** | 0[0, 0] |
| Midbrain peduncle contra | 0[0, 0] | **-1.05[-1.75, 0]** | 0[0, 0] | **0[0, 0.73]** |
| Central midbrain | **0.6[0, 1.97]** | 0[0, 0] | 0[0, 0] | 0[0, 0] |
| Pontine tegmentum | 0[0, 0] | **-0.05[-1.2, 4 0]** | 0[0 0] | **-0.86[-1.59 -0.21]** |
| **Diffuse injuries\*** |  |  |  |  |
| Volume | **-3.57[-4.9, -1.92]** | **-0.94[-1.74, -0.01]** | **-3.5[-4.53 -2.7]** | **-0.87[-1.54, -0.34]** |
| MLS | **-2.94[-4.41, -1.64]** |  | **-1.12[-1.75, 0.51]** |  |
| IVH | 0[0, 0] |  | **-0.58[-0.79, -0.41]** |  |

Data given as median [25%-IQR, 75%-IQR] of the weights obtained over the 500 cross validation iterations. Negative values indicate prediction of being unconscious, positive values of being conscious.
ROI: Region of Interest; ipsi: ipsilateral; contra: contralateral; MLS: Midline shift; ICH: Intracerebral Haemorrhage; IVH: Intraventricular Haemorrhage. \*: Volume weights correspond to 10ml units, MLS weights correspond to 1mm changes and IVH was dichotomized as present or absent.

Patients unconscious at time of MRI that were predicted to be conscious at ICU discharge more frequently were arousable than those that were predicted to remain unconscious, although this difference did not reach statistical difference (61% [N=25/41] vs 33% [N=4/12]; p-value = 0.1). In univariate analysis, GCS and FUNC scores were associated with level of consciousness at time of MRI and ICU discharge but primary ICH score was only associated with level of consciousness at ICU discharge (Table S2).

### Confounders

The model trained to predict consciousness at the time of MRI retained cumulative doses of midazolam and fentanyl together with the imaging parameters (Supplementary Material). However, performance of the models was not improved by including sedative doses (cumulative or current), cortical and cerebellar ROIs, or measures of metabolic disarray (i.e., renal insufficiency, glucose level).

### Functional outcome at 3 months

Three-month GOS-E were obtained for 92% (N=145/158) of the patients. Mean GOS-E was 4 (IQR 1, 6; Table 1), with 53% (N=77/145) of patients having a favourable outcome (GOS-E ≥ 4: 4% GOS-E 8, 19% GOS-E 7, 14% GOS-E 6, 1% GOS-E 5, 14% GOS-E 4), 19% (N=27/145) in a vegetative or totally dependent state (3% GOS-E 2, 15% GOS-E 3), and 28% of patients being dead (N=41/145; GOS-E 1). FUNC score reliably predicted functional independence (GOS-E ≥ 4) at month 3 (32% (N=6/19) with a FUNC score ≥ 9 vs 3% (N=1/30) for those with a FUNC score of < 9; p-value = 0.01). Patients that were unconscious at the time of MRI but predicted to be conscious at ICU discharge based on our model, more frequently were observed conscious at any time in the 30 days following the MRI (54% [N=22/41] vs 33% [N=4/12]; p-value = 0.3) and had better 3-month functional outcomes than those that were predicted to still be unconscious at discharge (GOS-E ≥ 4: 35% (N=13/37) vs 0% (N=0/12) vs; p-value =0.02; Figure 4).

## Discussion

More than half of ICH patients that are unconscious on admission are dead at one year after the bleed, and even though our ability to accurately prognosticate recovery is dismal, the primary mode of death is withdrawal of life sustaining therapies [24]. Accurate prediction of functional outcomes is challenging in the acute setting and confounded by biases contributing to the self-fulfilling prophecy of poor outcomes [25,26]. In this study we show that lesions identified on MRIs obtained within one week of ICH not only correlate with level of consciousness at the time of MRI but, more importantly, are able to identify patients that will recover consciousness prior to ICU discharge and have better 3-month functional outcomes. The identified predictors confirm previous findings (ICH volume, MLS and IVH) but also provide new insight on subcortical structures implicated in the physiology of consciousness [1–7, 12].

We confirm that measures reflecting the impact on both hemispheres (i.e., ICH and oedema volumes, MLS and IVH) are major determinants of impairment and recovery of consciousness, survival and functional outcomes. ICH volume is a well-known prognostic factor for both 30-day mortality and 90-day disability [15,16]. MLS is linked to high ICH and oedema volume and is clinically associated with herniation. Both large ICH volumes and MLS are seen in patients with increased intracranial pressure. All of these variables may cause bilateral impairment of widespread brain regions, which frequently is associated with unconsciousness [2].

Additionally, we identified three main subcortical structures implicated with consciousness impairment. These findings further support the role of the pontine tegmentum and midbrain peduncles, which have been implicated reliably in several clinical studies [1–3, 7]. Interestingly, the caudate nucleus, which has been implicated in wakefulness in the rat [27], is also included in many models of human consciousness as part of the frontal cortical–striatopallidal–thalamocortial loop systems [2,4,6]. According to the mesocirctuit theory, a decrease in the indirect excitatory activity of the Medium Spiny Neurons of the caudate and the putamen nuclei (special type of GABAergic inhibitory neurons) on the thalamus could explain an alteration of consciousness [6]. Hypometabolism of the caudate has been reported in unconscious patients [28] and, the caudate nucleus atrophies in chronic disorders of consciousness [8,29,30]. Reports of isolated bilateral caudate lesions are very rare but have been seen in patients with impairment of consciousness ranging from disorientation and confabulations to somnolence [31,32]. Caudate ICH has been previously associated with impairment of consciousness, however, since the caudate nucleus forms the wall of the lateral ventricle, it frequently is associated with IVH, hampered causal inference [33]. In our study, the majority of the patients with caudate lesions also had IVH. However, these patients were less likely to be conscious, both at time of MRI and ICU discharge, than patients with IVH in the absence of a caudate lesion (see supplementary material). At a minimum, caudate lesions appear to play a mediating effect on the relationship between IVH and impairment of consciousness. Finally, it is worth noting that the weighs attributed to caudate lesions were systematically greater than for IVH. In light of these findings, the present study provides further support implicating lesions of the caudate nucleus in impairment and early recovery of consciousness, independently from the frequently associated IVH.

The patient cohort studied here is not necessarily representative for all ICH patients as our enrolment bias analysis illustrates. Patients that were included tended to have slightly less neurological impairment, smaller haemorrhages, amyloid aetiology, and better outcomes. This is likely a reflection of provider safety concerns for MRI scanning and family preferences.

This study has several limitations. First, assessment of consciousness relied on a previously described, standardized neurological assessment [17] instead of a scale specifically developed for the assessment of consciousness such as the Coma Recovery Scale Revised (CRS-R) [34]. However, the CRS-R has some limitations in the ICU setting as it was primarily developed for patients in the subacute and chronic rehabilitation setting. Assessments with the CRS-R are time consuming posing a challenge in a hectic ICU environment during which patients consciousness level often fluctuates. We acknowledge that this assessment of consciousness will likely underestimate the presence of conscious and does not capture patients with cognitive motor dissociation [35,36]. Second, assessments of consciousness in patients with aphasia and delirium may be challenging. To capture nonverbal command following in aphasic patients we assessed, both verbal and non-verbal (i.e. mimicking) commands. Delirium in general and hypoactive delirium in particular are common in acutely brain injured patients and can interact with consciousness assessments. This confounder will affect any behavioural assessment in brain injured patients including the CRS-R [37]. However, the vast majority of patients with hyperactive delirium would be expected to demonstrate at least intermittent command following. Third, sedation is frequently used in the critical care setting and can confound assessments of consciousness. We minimized doses of sedation as recommended in guidelines [38] and systematically accounted for sedation given at the time of and preceding the assessment at time of MRI. Note that this limitation only applies the model trained at time of MRI since none of the patients received sedation at time of ICU discharge. Fourth, MRI based assessments of haemorrhage can be challenging as MRI signal changes are observed over time [39]. Subacute haemorrhages typically appear as hypointense signal in FLAIR sequences between 2-7 days, which was within the inclusion criterion in this study. Finally, confirmatory investigations to validate our findings on an independent dataset will be necessary in future studies.

Taken together, our results suggest that measures of injury obtained from routine clinical MRI sequences may allow to predict failure to recover consciousness by ICU discharge and functional outcomes in patients with acute brain injury more accurately. Focal lesions in key structures within previously described models of consciousness together with measures related to mass effect of the haemorrhage predict early recovery of consciousness. Both, adding a comprehensive assessment of structural connectivity between these key structures (i.e., using diffusion tensor imaging analysis) [40] as well as quantifying functional connectivity (using functional imaging or EEG markers) may further strengthen the accuracy of this model.

## Acknowledgement

We thank the nurses, attendings, fellows, and neurology and neurosurgery residents of the Neuroscience ICU for their overall support of this project. This publication was supported by the DANA foundation (JC) and the James S. McDonnell Foundation. BR received postdoctoral grants from “Amicale des Anciens Internes des Hôpitaux de Paris & Syndicat des Chefs de Cliniques et Assistants des Hôpitaux de Paris” (AAIHP - SCCAHP), “Assistance Publique – Hôpitaux de Paris” (AP-HP), and the Philippe Foundation. We are grateful to Dr Caroline Der Nigoghossian for her help with sedation data.

**Authors Contributions**
Study concept and design: BR, KD, ASR and JC. Data collection: ASR, KI, CC, AM and DR. Analysis and interpretation of data: BR, KD, ASR, BR, MM, CMM, GL, AE, VP and JC. Drafting of the manuscript: BR and LN. Critical revision of the manuscript for important intellectual content: BR, MM, SP, SA, CMM, GL, AE, VP, SC, AMB and JC. Statistical analysis: BR, KD, CMM, GL and JC. Study supervision: CC, AM and AV. BR, KD and JC had full access to all the data in the study and take responsibility for the integrity of the data and the accuracy of the data analysis.

